# Spatial-temporal dynamics of a microbial cooperative behavior robust to cheating

**DOI:** 10.1101/2020.01.23.914481

**Authors:** Hilary Monaco, Tiago Sereno, Kevin Liu, Caleb Reagor, Maxime Deforet, Joao B. Xavier

**Affiliations:** Weill Cornell Medicine, Tri-Institutional PhD Program in Computational Biology and Medicine, New York, NY, USA; Program for Computational and Systems Biology, Memorial Sloan-Kettering Cancer Center, New York, NY, USA; Summer Undergraduate Research Program, Memorial Sloan-Kettering Cancer Center, New York, NY USA; Sorbonne Université, CNRS, Institut de Biologie Paris-Seine (IBPS), Laboratoire Jean Perrin (LJP), F-75005, Paris, France

## Abstract

Individuals living in dense populations control their behaviors by sensing, integrating and responding to many cues. How can these processes enable the evolution and stability of cooperative behaviors that could easily be exploited by cheaters? Here we shed light on how bacteria regulate cooperation by studying swarming in *Pseudomonas aeruginosa*, a behavior requiring cooperative secretions of rhamnolipid surfactants to facilitate collective movement over surfaces. By combining fluorescent imaging and computational analyses we show, counterintuitively, that rhamnolipid expression peaks at swarming edges. We then show that the integration of competing diffusive cues—quorum sensing signals and growth-limiting nutrients—enables *P. aeruginosa* to communicate across centimeters and adopt expression patterns unseen in well-mixed liquid culture. Integration of multiple cues enables robustness against cheating even when we experimentally perturb the quorum sensing system. Taken together, these results illuminate how the integration of cues in spatially structured communities can stabilize cooperation.

## Introduction

Cooperation between cells allows microbes to contribute to multicellular communities ranging from antibiotic resistant biofilms (Costerton et al. 1999; Lee et al. 2010), fruiting bodies (Velicer and Vos 2009), swarming motility (Yan et al. 2019) and impact macroscopic organisms in ways that no individual microbe alone could (Singh et al. 2000; Rutherford and Bassler 2012). However, the microenvironments experienced by individual microbes living inside a densely microbial community are dynamic, densely packed and competitive (Granato et al. 2019). Cooperative traits can come at a cost to individuals because they require resources that could otherwise be used to grow (Griffin et al. 2004). How can microbial cooperative traits evolve and remain stable in nature in the competitive environment of bacterial communities? Understanding the cell-level computation that leads to evolutionary robustness of cooperative behaviors remains an open problem in sociomicrobiology. Spatial structure is key to the evolution of cooperation. Even a costly cooperative trait can be preserved as long as the benefits of cooperating can be localized to regions of highly related individuals (Nadell et al. 2010; Nadell et al. 2013; Kim et al. 2014; Drescher et al. 2014).

Microbes have the ability to sense, integrate and respond to diffusible queues by changing their gene expression. The concentrations of diffusible molecules that cells consume (such as nutrients) or produce (such as quorum sensing signals) change in time and with the dynamics of the rest of the surrounding cellular community. The spatial distribution of growth-limiting resources influences the development of structured populations, even in macroscopic communities: Vegetation growing in arid landscapes where water is the most limiting resource often show spatially structured patterning (Lejeune et al. 2004; Rietkerk et al. 2002; Rietkerk et al. 2004). The interactions between plants in arid environments can be described using an ecological kernel, where positive interactions represent nutrient preservation and negative ones, nutrient competition. Ecological kernels can also provide insights into the spatial structure of microbial communities (Deng et al. 2014), by describing how competing diffusible signals—growth limiting nutrients and quorum sensing signals—can influence the expression of a cooperative gene.

Here we investigate how the environment experienced by microbes influences their dynamics of cooperative behavior in spatially structured environments. We focus on the gram-negative *Pseudomonas aeruginosa*, an opportunistic pathogen often used as a model to study bacterial social behavior. *P. aeruginosa* builds thick antibiotic resistant biofilms that are life-threatening lung infections to cystic fibrosis patients and secrete a swath of disease-inducing virulence factors (Rutherford and Bassler 2012). These bacteria communicate by quorum sensing and transition between sessile (biofilm) and motile (swarming) life styles (Yan et al. 2019). *P. aeruginosa* swarms are of particular interest for their evolutionary robustness. This cooperative behavior allows a colony to grow over an order of magnitude larger in final population size (Xavier et al. 2011) but requires the expression of *rhlA* (Zhu and Rock 2008; Caiazza et al. 2005) and the subsequent secretion of massive amounts of rhamnolipid biosurfactant molecules (Caiazza et al. 2005; Déziel et al. 2003) that can amount to 20% of their dry mass (Xavier et al. 2011).

Secreted molecules required for swarming become publicly available once released, but the cooperative behavior remains robust to cheating. Wild type *P. aeruginosa* does not lose in competition against a Δ*rhlA* mutant unable to produce rhamnolipids thanks to the dynamic regulation of *rhlA* which integrates of quorum signals and information nutrient availability to delay expression to times when rhamnolipid secretion becomes affordable (Xavier et al. 2011; de Vargas Roditi et al. 2013; Boyle et al. 2015). The ability to regulate investment in a cooperative trait to avoid a fitness cost is termed Metabolic Prudence (Xavier et al. 2011), a strategy that may regulate many bacterial social traits (Xavier et al. 2011; Mellbye and Schuster 2014; Smith and Schuster 2019).

Rhamnolipids are a high carbon-content compound. Experiments tracking gene expression in liquid culture showed that if cells run out of carbon source, they shut off *rhlA* expression. When cells run out of either nitrogen or iron instead, cells ramp up *rhlA* expression and allocate carbon towards rhamnolipid synthesis. This is presumably to facilitate movement to more nutrient rich locations at no fitness cost. Adding quorum signals to the medium also amplifies *rhlA* gene expression in liquid culture, particularly when the cells are in early stationary phase (Boyle et al. 2015).

The regulation of *rhlAB* expression integrates nutrient and quorum signal information and depends on at least three diffusible molecules: a growth-limiting nutrient, and the hierarchical quorum sensing structure involving the quorum signal molecules 3-oxo-C12-HSL and C4-HSL (Latifi et al. 1996; Pearson et al. 1997; Ochsner and Reiser 1995; Ochsner et al. 1994; Wagner et al. 2003; Medina et al. 2003). According to the literature, all three diffusive inputs, the two auto-inducers as well as any small molecule growth-limiting nutrient, act on similar length/time scales. In addition, the ratio of the diffusion coefficients and decay rates for these molecules in bacterial growth media (Cornforth et al. 2014) indicate that the quorum signals could achieve high enough levels to reach and influence biomass that is multiple millimeters away. Still, the diffusible species may compete in their control of gene expression: growth nutrients such as nitrogen and iron should downregulate *rhlAB* whereas quorum sensing signals should upregulate *rhlAB*. The regulation of *rhlAB* in a spatially structured system may be quite complex and sensitive to environmental fluctuation.

Considering the initial seeding of a swarming assay, at the center of an agar plate, we hypothesized that the nutrient environment would deplete in the center of the swarm first and then proceed outward, standard to population motility theory, with the region of active growth localized to the edge of the swarming tendril at the interface between the population and growth limiting resources (Deforet et al. 2019). We expected quorum signals to follow the reverse pattern, building first in the center of the swarm with lowest levels at the swarming tendril tips. Given liquid culture literature, this lead to a hypothesis where rhamnolipids are largely being produced at the center of a swarm, where quorum signals are high and growth rate is low, with minimal production at the tendril tips where quorum signals are low and growth rate is high.

Here we analyze image timeseries of *P. aeruginosa* swarms fluorescently labeled for biomass production and P*rhlAB* activity (Supplementary Figures 1-3, and 5). We find that, contrary to our hypotheses, the edges of swarming tendrils emerge as the regions of highest cooperative gene expression. To interrogate the role of the diffusive inputs on *rhlAB* expression in describing this phenotype, we used immotile colonies seeded as in the classic Colony Forming Unit (CFU) assay to investigate how the interactions between colonies affect rhamnolipid production. Using these data, we fit an ecological kernel through regularized regression motivated by reaction-diffusion principles showing that both growth rate information and colony neighborhood configuration are critical to explain the complex gene expression we observed. Finally, we show that quorum signals, known to facilitate cellular communication over micrometer distances (Darch et al. 2018; Connell et al. 2010; Connell et al. 2014), are capable of centimeter-scale communication between *P. aeruginosa* colonies. Further, while perturbation by quorum signals in liquid culture never showed significant alteration to *rhlAB* expression, the same perturbation in the spatially structured system surprisingly revealed an over-expression phenotype that was nonetheless cost-less in both bacterial colonies and motile swarms. Taken together, these data show that there are new regimes of bacterial gene expression yet to be unlocked in spatially-structured systems. Our findings reveal new scales of bacterial communication and new dimension to the evolutionary robustness of bacterial cooperation.

## Results

### Expression of *rhlAB* peaks at the edge of swarming colonies

Competition for nutrient and quorum sensing are two types of cell-cell interactions mediated by diffusible processes that affect *rhlAB* expression. Their competing influences make the dynamics of *rhlAB* expression in motile swarms difficult to predict. We constructed a fluorescent imager inside an incubator to track cell growth and *rhlAB* expression directly in colonies on Petri dishes (Supplemental Figure 1), using a *P. aeruginosa* PA14 strain with a dual-label construct harboring PBad-DsRed(EC2) (Pfleger et al. 2005) driven by L-arabinose which was included in the plate media (Newman and Fuqua 1999) and P*rhlAB* -GFP (van Ditmarsch and Xavier 2011; Boyle et al. 2015). The constitutive expression of DsRed provided an indication of the local density of bacteria (Supplemental Figure 2), and the dynamical expression of GFP reported on the expression of *rhlAB*. The data was corrected for uneven lighting of the samples (Supplemental Figures 1, 3, see Methods).

Using this imaging device we investigated swarming (Figure 1a). Counter to our expectations, that *rhlAB* expression peaked at the tip of each swarming tendril rather than at the center of the swarming colony. To confirm this observation, we quantified expression along the length of three tendrils from four independent swarming colonies throughout the time course of the swarm (Figure 1b). The *rhlAB* expression increased with the distance from the swarm center in all cases. This dynamic coincides with an unexpected growth phenotype in the swarming tendrils (as reported by the red signal): The biomass across the entire tendril showed an exponential growth rate (Figure 1c inset), which was particularly surprising as the average spreading velocity of the tendril is linear at 3.2mm/h with a standard deviation of 0.8 mm/h.

**Figure 1:**
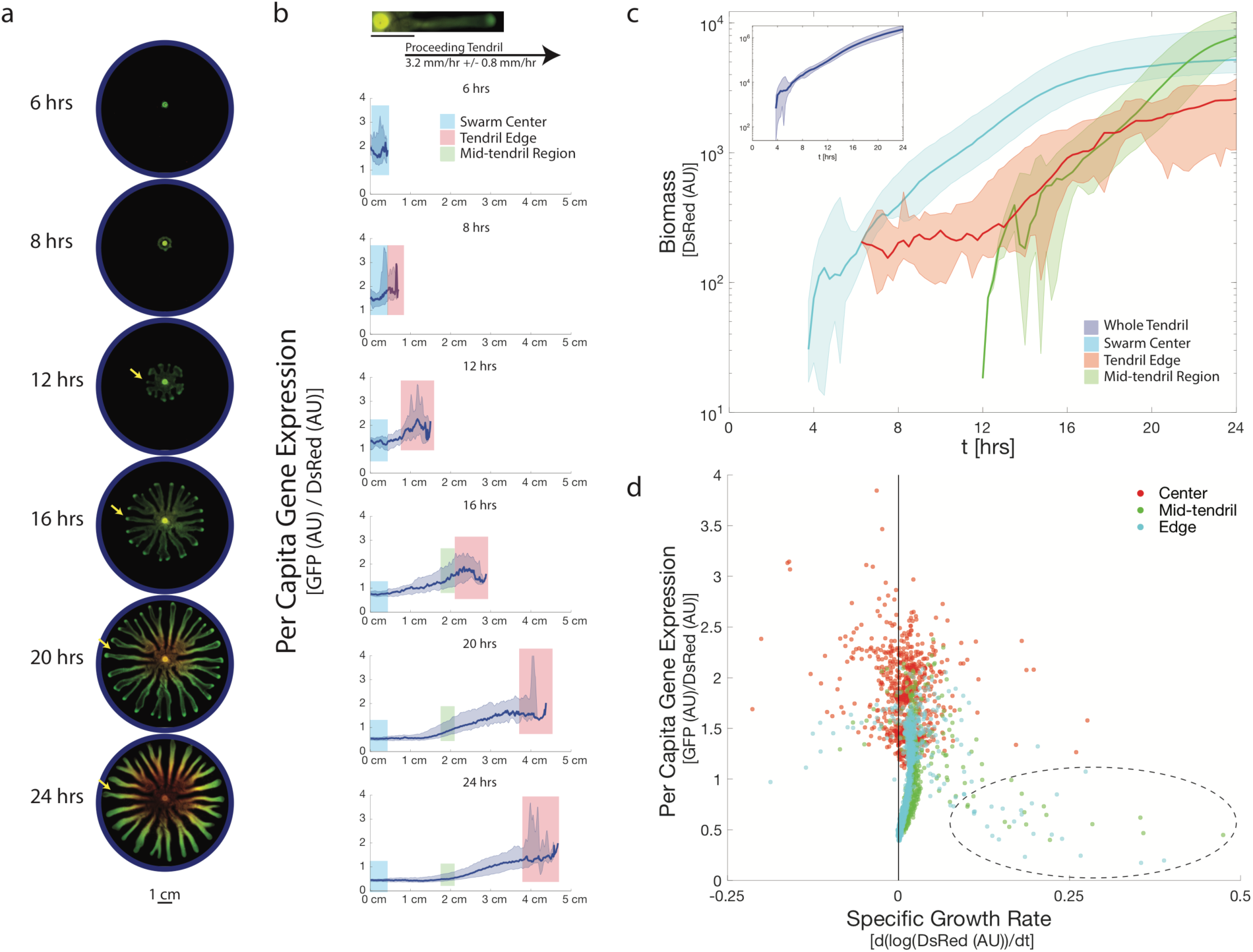
Swarming tendrils move with linear velocity, grow exponentially and show spatially segregated gene expression for rhamnolipid production. **a.** Swarms were imaged in 5-minute intervals across a 24-hour period. Cells are fluorescently labeled for biomass (constitutive marker for biomass PBad-DsRed (EC2) induced by L-arabinose in the plate media) (red channel) and activity of the *rhlAB* operon (through the promoter fusion P*rhlAB*-GFP) (green channel). Images of selected timepoints were background corrected (see Methods) and are contrasted to the maximum intensity found in all images in the timeseries. **b.** Per capita gene expression is tracked by the ratio of GFP (P*rhlAB*-GFP) to DsRed (used as a biomass marker) per pixel. Data across three tendrils in each of four independent swarms were combined for analysis. We observe that the highest investment is surprisingly found at the edge of the swarm. Median gene expression data is highlighted and full range of the data is shaded. Max and min data was smoothed again for visualization. All tendrils were aligned to start at the same location for analysis though tendrils reach different final lengths. **c. Inset:** Total biomass identified along the length of isolated tendrils with time. We observe that the whole tendril appears to sustain an exponential rate of growth throughout the timeseries. **Main Panel:** Biomass with time for three sections of the swarm: Center (blue), mid-tendril (cyan) and swarm edge (red) for the same tendrils as in **b**. Data normalized for size of region isolated. Median data plotted in bold, area shaded is full range of the data. Max and min data was smoothed for visualization. Note that the mid-tendril maintains an exponential growth rate after the edge of the tendril has passed. We also find that the edge of a swarming colony appears to sustain a rate of increase in localized biomass similar to the mid-tendril. **d.** Per capita gene expression plotted against the growth rate of each region of the swarm. Growth rate determined as the derivative of the log of the red data. We find the swarm center and mid-tendril regions have lower per capita gene expression at high growth rate and higher at low growth rate, consistent with previous reports (Boyle et al. 2015; Xavier et al. 2011). We find that at our lowest observed growth rates, gene expression drops, consistent with (Boyle et al. 2015). Per capita gene expression at the swarm edge is noticeably higher than the mid tendril or swarm center, though the variation is not clearly explained by the growth rate of biomass localized to the swarm tip.

To characterize how an exponential growth rate could emerge in a tendril advancing linearly, we analyzed three regions within the swarming tendril: the center of the swarm, a fixed-sized region in the middle of the tendril, and the edge. Pixels were isolated and used to calculate the average behavior in each region (Figure 1c). These swarms formed tendrils after an average of 7.25 hours of growth with a standard deviation of 21 minutes. Surprisingly, the biomass level at the center of a swarm continued to grow exponentially long after tendrils had formed and started to move away from the initial seeding location. Similarly, the mid-tendril region also maintained an exponential growth phase both regions showing doubling times of approximately two hours.

The edge of a swarming colony is where fresh nutrients abound and where growth is presumed to be fastest. However, part of the biomass produced is left behind as the edge of the tendril as moves forward in a traveling wave (Deforet et al. 2019), and that biomass seeds the mid tendril. Our observation that the mid tendril maintains exponential growth indicates that the edge moves forward before the nutrients below are fully consumed.

As we do not control for flux of biomass between tendril regions, the growth rate measured represents a combination of growth and migration into and out of each region. However, as the general flow of the biomass is away from the swarm center, to first approximation the flux into the center or mid tendril can be neglected. The growth rate calculated for the edge of a tendril likely underestimates the cellular growth rate in that region because there is unlikely to be flux into the tendril tip, and only flux out of it (the biomass left behind as the edge moves away from the seeding location).

To investigate whether any region of the swarm behaved similarly to expected metabolically prudent dynamics, we analyzed each region’s per capita gene expression with respect to the corresponding growth rate (Figure 1d). In the center and mid-tendril of the swarm, we observe that high growth rates correlate with lower levels of per capita gene expression (Figure 1d dashed circle). This indicates that the cells may indeed be titrating gene expression in accordance with local nutrient availability as in liquid culture (Boyle et al. 2015). However, at the edge we find no correlation between *rhlAB* expression and measured growth rate. As we are likely underestimating the true growth rate of the biomass at the edge, this result was puzzling given current knowledge of the inherent gene expression control. Overall, locally the swarm tendril behind the swarm tip seemed to be behaving in accordance with known metabolic prudence, however, globally the high per captia gene expression localization to the tendril tip remained unexplained.

### Experiments on hard agar recapitulate spatial-temporal dynamics of *rhlAB* expression without the complication of movement

Studying the dynamics of growth and *rhlAB* expression in swarming colonies is complicated by the difficulty of separating growth rates from motility flux in the regions examined above. Swarming requires an agar concentration of 0.5%, and increasing the concentration to 1.5% is enough to prevent swarming (Xavier et al. 2011). In hard agar, a strip of immotile PA14 (Supplementary Figure 4a) revealed the same pattern of *rhlAB* expression in this immotile model of a tendril with *rhlAB* expression peaking at the edges (Supplemental Figure 4c). However, the immotile tendril was unable to sustain an exponential growth rate (Supplemental Figure 4b), indicating that this gene expression phenotype may be able to be explained by the diffusive inputs to this system.

To probe the role of solute diffusion on *rhlAB* expression we continued to use hard agar experiments to prevent motility (Figure 2a). Using extreme dilution of bacterial inocula (Figure 2b) we seeded colony forming units (CFUs) and tracked the development of those colonies from single cells to mature colonies. We computed growth and *rhlAB* expression for each individual colony (Supplemental Figure 3, 5). Using a range of experiments, we varied the number of colonies and their local distribution in each Petri dish to produce thousands of growth and *rhlAB* expression curves and capture a wide diversity of gene expression behaviors.

**Figure 2:**
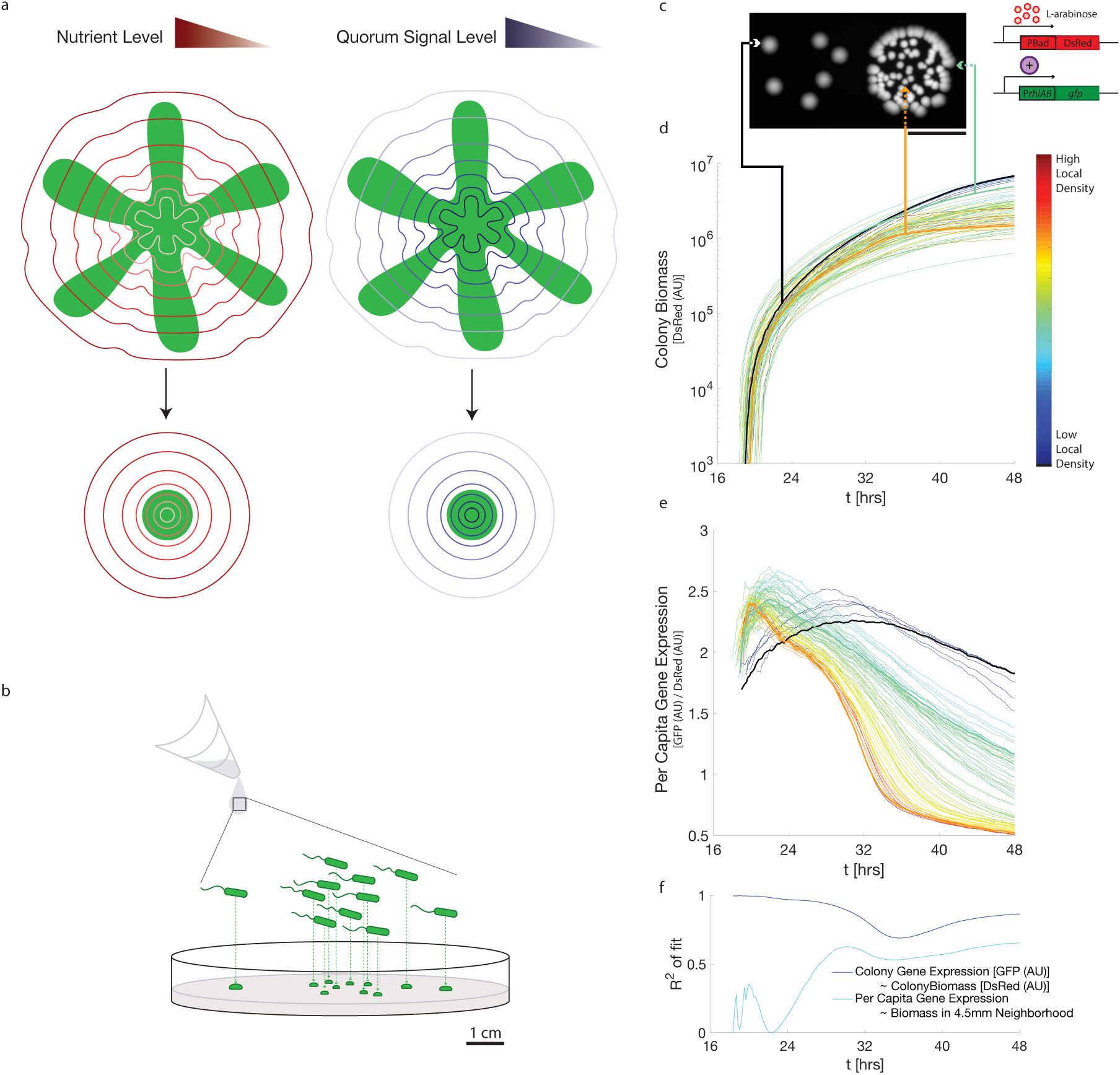
Colony Forming Units as a model system for emergent growth and cooperation patterning. **a.** Growth and cooperation dynamics that emerge from nutrient and quorum signal diffusion can be studied in hard agar diffusive systems. [Top] Cartoon depicting hypothesized nutrient and quorum signal fields in developed swarms. [Bottom] Cartoon depicting the simplified fields predicted of developing colonies. **b.** Method of generation for of Colony Forming Unit (CFU) plates. A washed and dilute culture is placed and left to dry on the plate. Colonies that emerge were seeded from individual single cells. (See Methods) **c.** CFU image at 48 hours. Data characterizes colony biomass as indicated by intensity of the shown DsRed image. Scale bar 1 cm. Colonies indicated are highlighted in D and E. Schematic describes fluorescent labeling used. (See Figure 4a for more information on the *rhlAB* regulation pathway.) **d.** Analysis of image timeseries (see Methods and Supplementary Figure 1-3 and 5) generates a growth curve for each colony. Colony growth trajectories are colored by the number of colonies within a 4.5mm radius of the focal colony. Variation in colony growth curves correlates, as expected, with spatial configuration. Colonies highlighted demonstrate the variation observed in the dataset. **e.** Colony per capita gene expression of *rhlAB* operon calculated as in Figure 1. Coloration of the expression data as in D reveals that in this dataset, many neighbors in a 4.5mm radius correlates with an earlier peak in per capita gene expression. **f.** Explanation of variation observed in colony per capita gene expression. Data at each timepoint is independently fitted to the indicated model and goodness of fit shown. We observe that much of the variation in colony gene expression varies with colony biomass (Dark blue curve). However, this correlation decays with time. At later timepoints, variation in colony per capita gene expression, can be explained by the amount of non-self biomass in a 4.5mm neighborhood around the focal colony. This indicates that the colonies may be able to influence the gene expression of one another.

Our data showed that colonies located in regions of higher local density grew to smaller colony sizes at 48 hours compared to colonies in less dense regions of the same plate. This is expected from the effects of nutrient depletion from local crowding (Figure 2d) and can be captured by the variation in the amount of biomass within a 4.5mm radius neighborhood of each colony. The *rhlAB* expression curves, however, revealed an unexpected diversity of dynamics that showed a very complex dependency on local neighborhood. We found that while the peak per capita gene expression in a focal colony correlated with the amount of biomass within a 4.5mm radius neighborhood, but the correlation between a colony’s neighborhood its per capita gene expression varies in both amplitude and sign with time (Figure 2e).

One way to characterize these data was to ascertain how the variation in each colony’s per capita gene expression could be explained by the current state of the system. We hypothesized that if the *rhlAB* expression level of each colony depended only on the focal colony itself then each colony’s *rhlAB* expression signal would correlate with its corresponding biomass signal. Conversely, if colony-colony interaction played a key role then *rhlAB* expression would not correlate with the colony’s biomass level. To test this, we took the *rhlAB* gene expression at each timepoint and asked if the variation could be explained by the size of each colony at the same timepoint. We found that early in our timeseries, *rhlAB* expression does correlate well with red levels (Figure 2f – Dark Blue Curve). However, later in the timeseries, there is a drop in the *rhlAB* expression variation that can be explained by the biomass signal. To investigate the remaining variation we calculated the per capita gene expression by calculating the ratio of the total green fluorescence to the total red fluorescence for each colony. We find that as the correlation between the GFP and DsRed signal declines the variation in the ratio that can be explained by the biomass in a focal colony’s 4.5mm neighborhood starts to increase (Figure 2f – Cyan curve). This analysis indicated that colony-colony interactions may indeed alter the *rhlAB* expression of each individual colony.

### Growth state alignment and regularized regression quantifies the interaction between neighboring colonies

Next we sought to detail the function by which each radius of a focal colony’s neighborhood influenced its *rhlAB* expression. We collected timeseries from colonies started from single cells, generating thousands of independent experiments in high throughput. Each colony came above detection by our pipeline at a slightly different time, even among colonies on the same plate. This made it difficult to understand how two colonies with similar growth trajectories (Figure 3b inset) could have such different per capita gene expression patterns (Figure 3b main panel). However, we noticed that the growth rate of each colony was predictive of its per capita expression dynamics (Figure 3b main panel). At high growth rate, colonies had low levels of per capita gene expression. Between a small range of low growth rate, the per capita gene expression peaked. Below a threshold growth rate, gene expression turned off. Using this information, we grouped colony data by growth rate. These similarities allowed us to investigate how the neighborhood surrounding a colony could generate variation in per capita gene expression.

**Figure 3:**
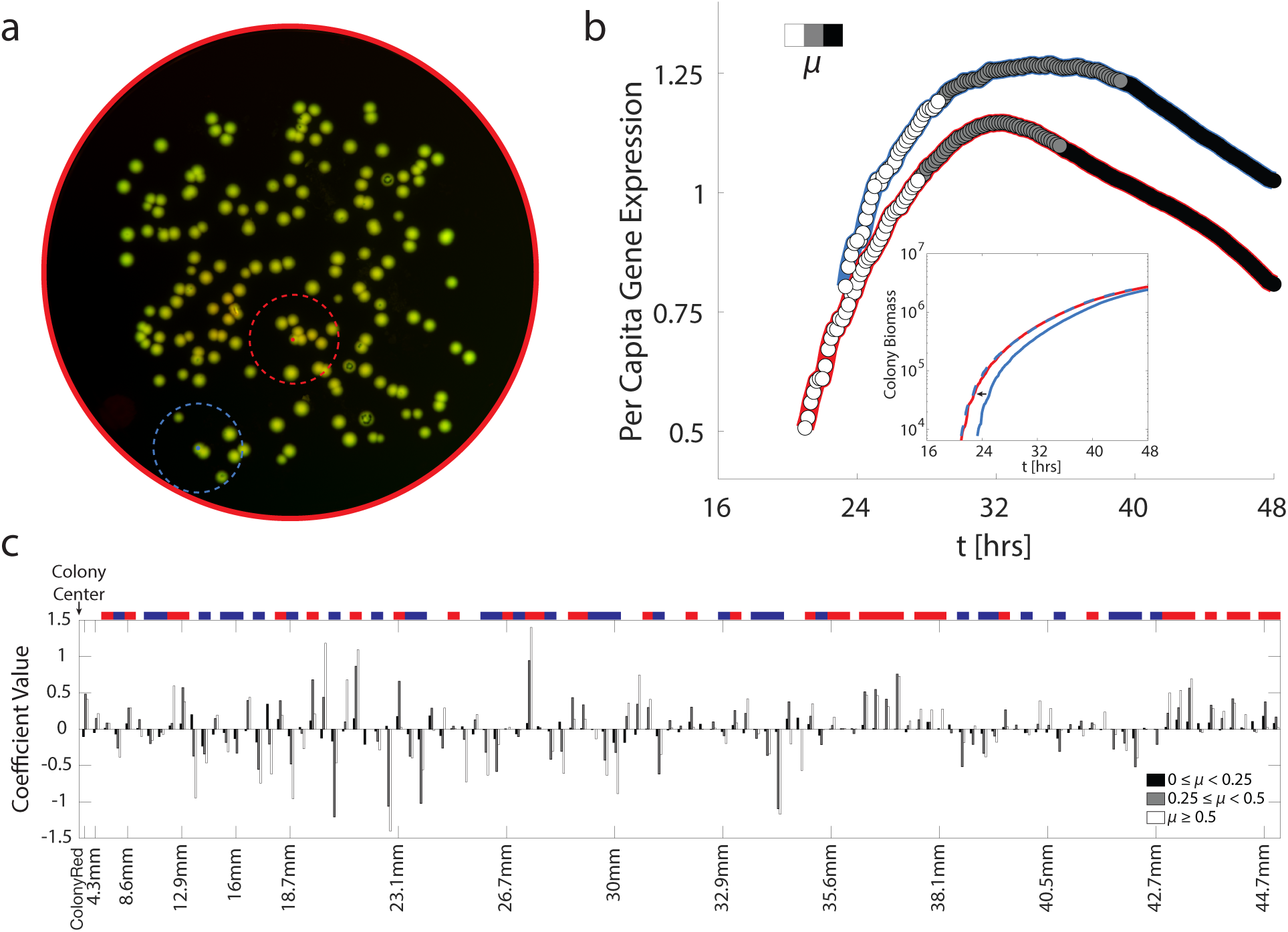
The spatial-temporal gene expression patterning from the integration of nutrient and quorum signal information can be described through a spatial kernel that changes with local density. **a.** Two colonies and respective example neighborhoods, indicated by blue and red dashed circles. **b.** Colonies with similar growth curves can show strong differences in gene expression dynamics but similarities appear when data is grouped by growth rate. [Inset] Two colonies (the ones indicated in **a**) have similar growth dynamics, as shown by the time shifted blue colony curve (dashed blue line). [Main Panel] Per capita gene expression patterns vary between the two colonies, however growth rate, *μ*, (high (white), medium (gray), low (black)) correlates with changes in gene expression dynamics. Periods of high growth rate correlate with increasing levels of gene expression, intermediate growth rates correlate with peak levels of gene expression, and low growth rate correlates with declining per capita gene expression. **c.** Ridge regularized results describing the kernel of interaction in each of the three growth rate bins. Data across four experiments with configurations like that shown in **a** are included. In general, the coefficient describing the influence of biomass in that neighborhood on a focal colony is either positive (red) or negative (blue) with the scale of the response varying with the growth rate of the focal colony. Coloration above the plot is used whenever all three growth rate bins show the same coefficient sign, inclusive of 0. R^2^ values in Supplementary Table 1.

To quantify the colony-colony interaction, we inferred a spatial interaction kernel directly from our data. We computed the amount of biomass (using the red signal) in concentric neighborhoods around each focal colony (Figure 3a). These neighborhoods were then used as features to explain variation in *rhlAB* per capita expression. We grouped data across growth curves and four independent experiments with similar configurations of colonies by growth rate (Figure 3b) and we applied ridge regularization to fit three models, one for each growth rate bin (Figure 3c). The results revealed a complex spatial-temporal pattern of activation and inhibition that results from the different length-scales of the diffusional factors as the colonies developed. However, we noticed the coefficients corresponding to the influence of each neighborhood on a focal colony’s behavior showed a pattern: The coefficients of a given neighborhood often shared the same sign across all three regressions, but differed in amplitude (Figure 3c blue white and red bar). This may indicate that in a given configuration, there is a fixed spatial interaction kernel and the colony’s growth rate is indicative of the colony’s ability to respond to the information in that kernel.

### Signal-negative mutants validate distance-dependent activation of *rhlAB* expression

Next, we sought to confirm that one of the inputs responsible for the rich dynamics in *rhlAB* expression was a response to quorum signals. To do this, we utilized a mutant unable to produce the 3-oxo-C12-HSL and C4-HSL signals, PA14 Δ*lasI*Δ*rhlI* double-labeled in the same way as the WT, as a quorum signal receiver (Figure 4a). To isolate the quorum signal response, we focused on a colony’s response to the C4-HSL molecule, the furthest downstream of the two quorum signals. Tracking P_*rhlAB*_ activity showed whether that colony had sensed both quorum signals. We added 1µM 3-oxo-C12-HSL to the plate media and placed 4µL of 5µM C4-HSL on a filter paper on the center of a petri dish and we tracked the growth and *rhlAB* expression in colonies started from single cells seeded around the filter paper (Figure 4b). The mutant colonies showed maximal *rhlAB* per capita gene expression at the colony peak that was inversely proportional to the colony’s distance to the filter paper (Figure 4c, d), R^2^ = 0.37. However, there was a batch effect that corresponded with the number of colonies on the plate where plates with fewer colonies (light blue data points) showed higher per capita gene expression overall than colonies with a denser colony seeding (dark blue and black data points).

**Figure 4:**
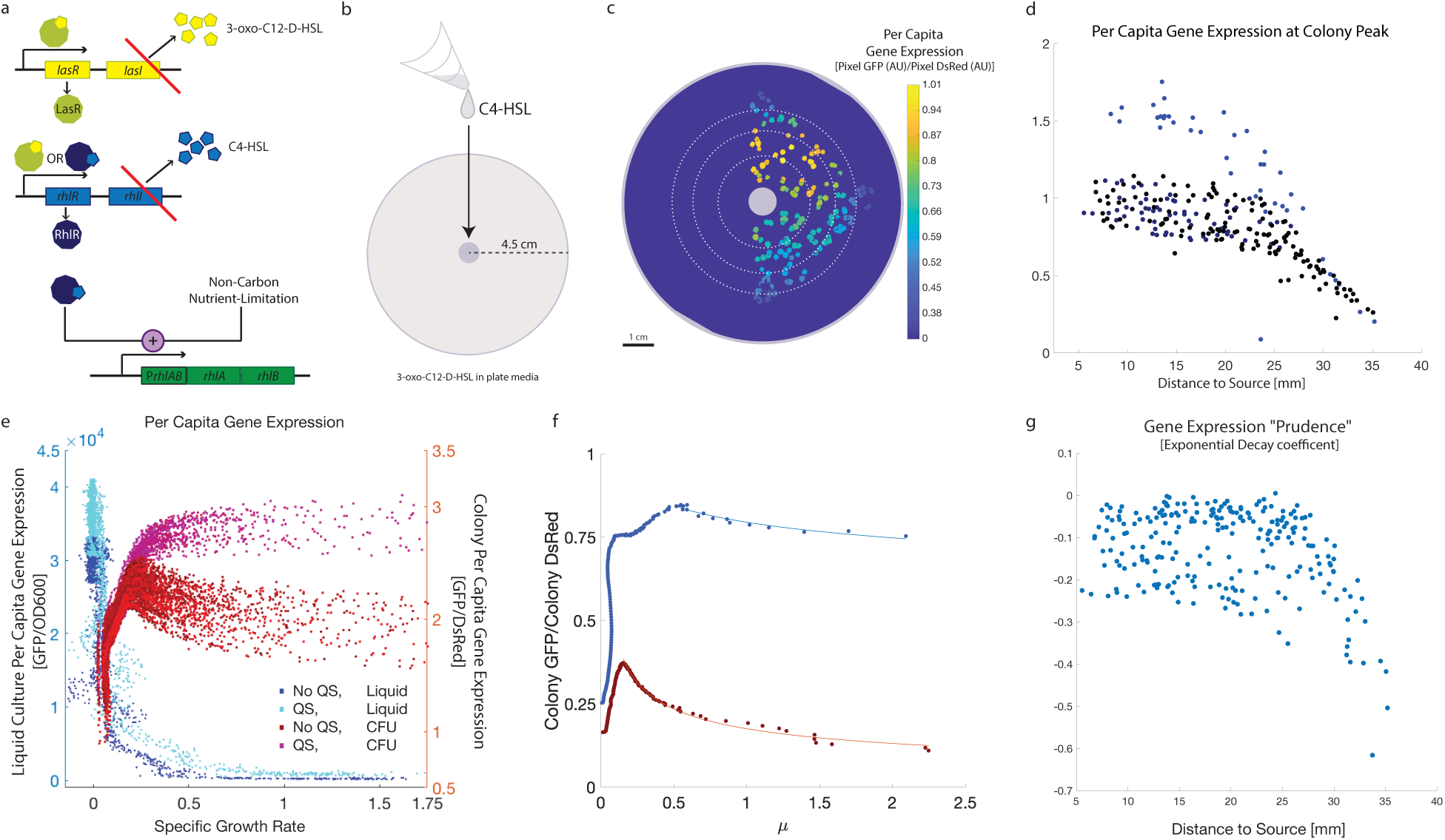
Perturbation with quorum signals reveals a spatially-linked expression pattern that scales with distance to the quorum signal source. **a.** The native molecular circuit determining *rhlA* expression. The molecular circuit and alterations (red lines) describes a signal mute quorum signal mutant that is unable to produce the two autoinducer molecules required for *rhlA* production. **b.** Experimental design to demonstrate the centimeter length scale of quorum signal response. The upstream quorum signal (3-oxo-C12-HSL) is added directly to the plate media and the downstream signal, C4-HSL, is loaded on a filter paper in the center of the plate. Colonies have been seeded around the filter paper and will respond if the colony observes both quorum signals. **c.** Signal mute colonies respond to diffusible quorum signals. Colony borders indicative of colony area after 24 hours of growth. Coloration indicates the maximum per capita gene expression achieved throughout the time-course at the colony peak. **d.** Per capita gene expression at colony peak correlates with the distance of the colony to the center of the filter paper. Data from three independent biological replicates with 40 (blue), 69 (dark blue) and 139 (black) colonies on the plates respectively. R^2^ = 0.3744. **e.** Liquid and spatially structured expression patterns with respect to measured growth rate. Liquid culture per capita *rhlA* gene expression measured as GFP/OD600 (Xavier et al. 2011) when grown in liquid culture with (Cyan) and without (Blue) exogenous quorum signals. There is no clear difference in the per capita gene expression (Blue Axis) with quorum signal perturbation in liquid. However, immotile colonies grown without quorum signals (red) and with quorum signals in the plate media (magenta) show a clear difference in per capita gene expression (Orange Axis). Growth rate is calculated as the derivative of the log(DsRed) data with respect to time. Data for each experimental configuration includes three biological replicates. CFU data was taken from plates where colonies were grown at low numbers in low local density to minimize colony-colony interactions. Each datapoint represents a time interval in a liquid or immotile colony timeseries. **f.** Signal mute mutants closer to the quorum signal source show a higher per capita gene expression regardless of growth rate. These dynamics are shown for two example colonies here. Colonies are indicated in **c**. To characterize the turn on pattern of this gene expression with respect to growth rate, we fit an exponential decay curve from high growth rate, to the growth rate with maximal per captia gene expression. The orange colony at 35.06 mm from the quorum signal source shows a strong correlation between per capita gene expression and growth rate. By contrast, the blue colony at 7.17 mm from the quorum signal source shows a response that is roughly independent of the growth rate and has a much smaller in magnitude exponential decay coefficient. **g.** Colony location relative to the quorum signal source (the filter paper) explains variation in temporal dynamics of per captia gene expression. Colony gene expression is characterized by the coefficient of the exponential decay fit of the data from maximal growth rate to maximal investment across the time course. This analysis reveals that colonies less than 2.5-3cm from the quorum signal source have low exponential decay coefficients indicating that the per capita expression remains close to the colony’s maximum value even during periods of high growth. Exponential decay coefficients beyond 25mm from the source vary linearly with their distance to the source with R^2^ = 0.60.

### Perturbation with quorum signals reveals a spatially-linked expression pattern that scales with distance to the quorum signal source

These experiments carried out with the signal negative mutant confirm that diffusible quorum signals explain part of the spatial-temporal pattern of *rhlAB* expression. We did not expect, however, that adding quorum signals to the media would influence the colony behavior of WT bacteria. Previous work done in liquid culture showed no change in total rhamnolipid production in WT bacteria grown with quorum signals added to the media (Xavier et al. 2011). Surprisingly, when quorum signals were added to the same plate media recipe, WT colonies seeded far from each other expressed more *rhlAB* during periods of higher growth rate, even when compared to other wild type (WT) colonies grown in similar configurations without added quorum signals (Figure 4e).

To understand the discrepancy between the liquid culture versus spatially-structured colonies, we looked to see if these expression dynamics replicated in the signal mutant. We found that colonies close to the filter paper expressed *rhlAB* more at high growth rate than those far from the filter paper source (Figure 4g). This behavior could be quantified by fitting colony expression at high growth rate with a decaying exponential function (Figure 4f,g). In doing so we uncovered a threshold-like detection response (Figure 4g). Colonies less than 2.5-3cm from the quorum signal source show a similar induction pattern with little variation in per capita gene expression at high growth rate, just as we observed in our WT colonies with supplemented quorum signals. Farther than 2.5cm away, the exponential decay coefficients vary linearly with colony distance to the source (R^2^ = 0.60) and have a very low peak per capita gene expression.

### Swarming is robust to cheating despite *rhlAB* overexpression by extrinsic quorum signals

To test whether this alteration in gene expression carried a cost to cell growth, we measured colony fitness in three independent ways. First, we looked for a change in the distribution of growth rates of the colonies in the first time interval after detection. A lower growth rate under quorum signal perturbation would have indicated a growth cost prior to detection. We found no difference between these growth rate distributions in experiments with and without quorum signal perturbation with colonies in similar configurations nor in plates with an alternative configuration also perturbed with quorum signals (Figure 5a). Next, we looked for a difference in the final colony size. If the colonies in the perturbation were smaller at the end of the timeseries, a growth cost may have occurred in a less obvious way during the time interval of colony observation. We found instead that the colonies that were grown with added quorum signals were the same size as colonies grown without quorum signal when in a similar configuration (Figure 5b). Finally, we looked for a transient cost, a temporal element to the behavior that could indicate a comparatively different state of growth when comparing datasets grown with and without quorum signals. We compared the distributions of the times when the colonies, come above detection. Here, we saw indeed that colonies grown with quorum signals can come above detection later than colonies grown without (Figure 5c). When quorum signal mutants are subjected to quorum signals they can stay in lag phase longer (Boyle et al. 2015), and this may be what occurred here for the for WT bacteria growing in media with added quorum signals.

**Figure 5:**
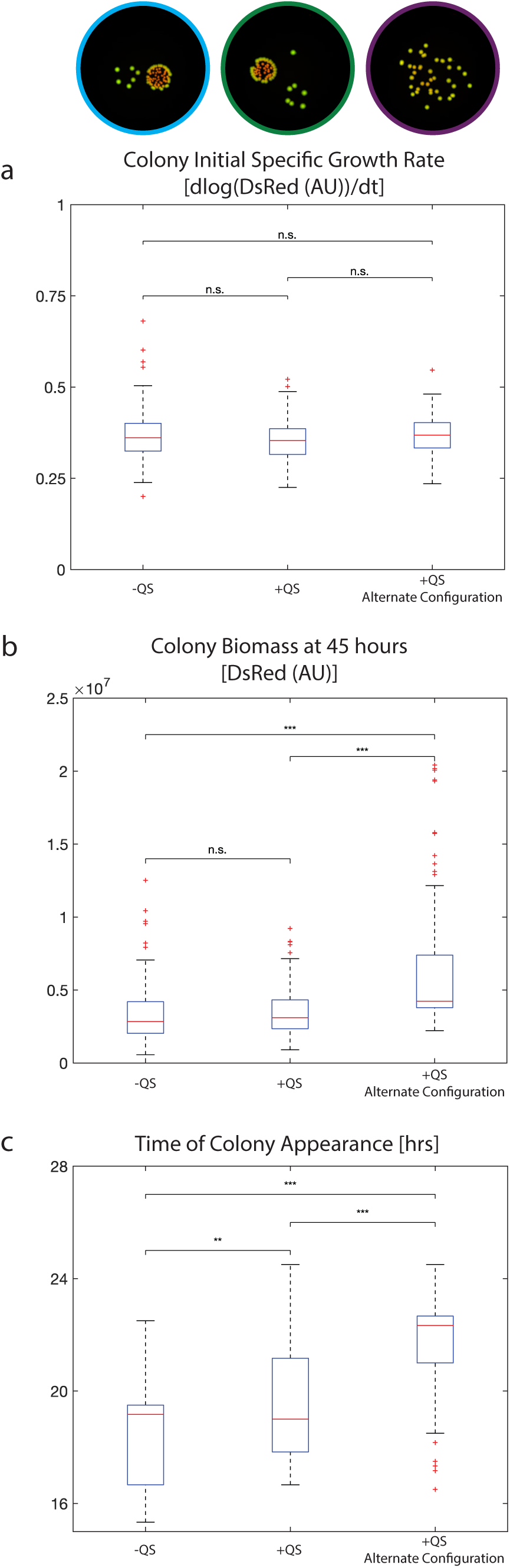
Quorum signal perturbation reveals no growth impact. **a.** Comparison of growth rate at the time of colony appearance with and without quorum signals in the media. Three biological replicates of three colony configurations were compared. Representative datasets pictured at the top. [Left configuration] colonies organized with high configuration variation. [Center configuration] colonies organized as in the left configuration, colonies perturbed with quorum signals in the media. [Right configuration] Colonies grown with low configuration variation, spread far apart on the plate. Media contains quorum signals at the same levels as the center configuration data. Colonies growth with or without quorum signals in the plate media show no difference in initial growth rate. Colonies grown at lower local density with quorum signals in plate media also show no difference in initial growth rate distribution. Colony identification algorithm was consistent across datasets [See Methods]. Significance measured by the Mann-Whitney test (See Supplemental Table 2). **b.** Comparison of final colony biomass at 45 hours with the same 9 datasets as in A. Colonies grown with quorum signals in the media were found to have a similar size distribution to colonies grown without quorum signals in a similar colony configuration by the Mann-Whitney test. Colonies grown farther apart grow to larger final size. Quorum signal perturbation does not lead to any apparent growth cost we observe at 45 hours. **c.** Comparison of time of colony appearance. Colonies grown without quorum signals are found to appear earlier than colonies grown with quorum signal. Significance measured by Mann-Whitney test.

To see if this phenotype also played a role in the motile swarming model system, we searched for an increase in *rhlAB* expression in the motile swarms. Indeed, we found that adding both autoinducers to the media media accelerated the onset of swarming by ∼30 minutes compared to swarms without supplemented autoinducers (Figure 6a).

**Figure 6:**
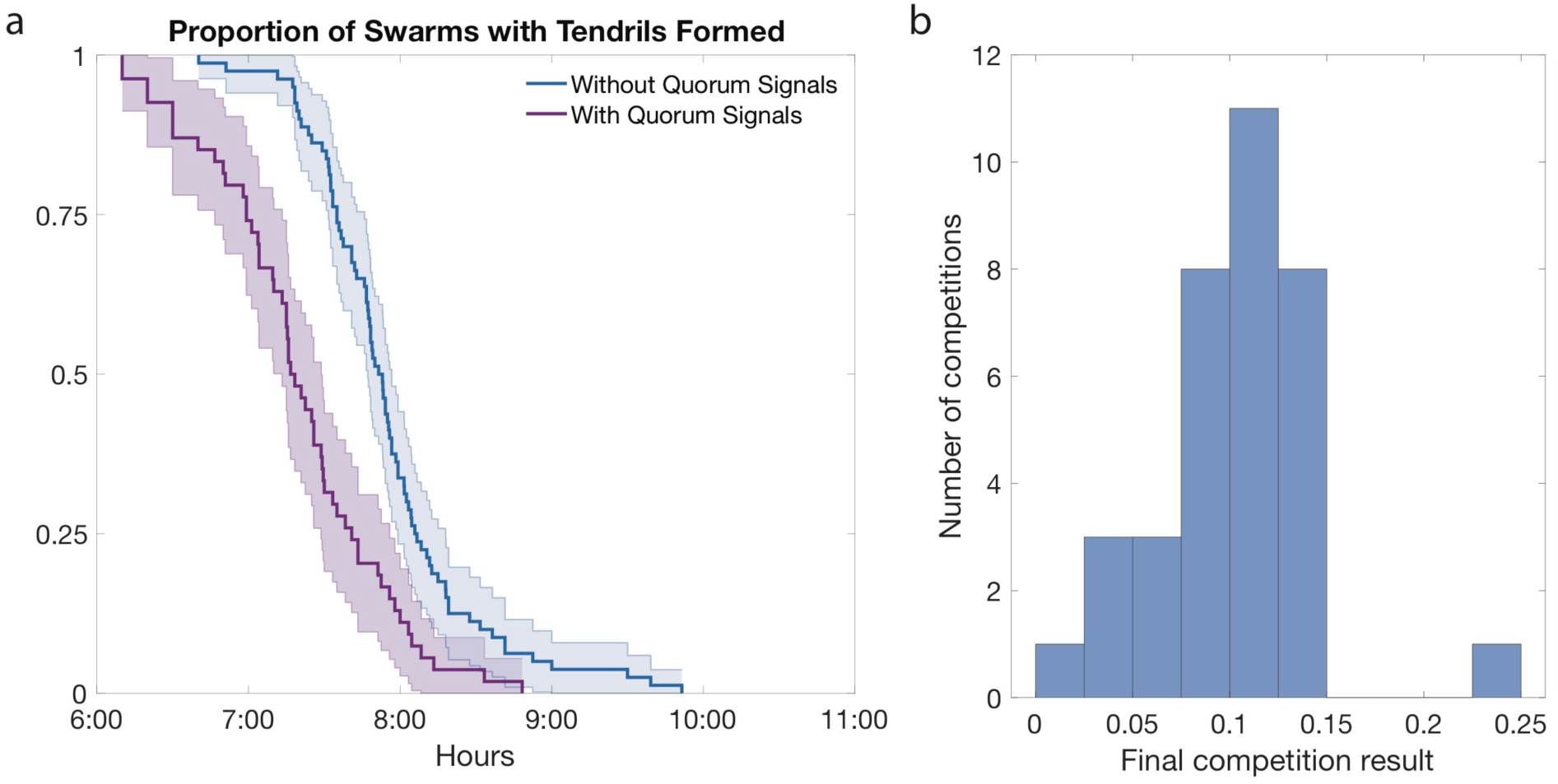
Quorum Signal perturbation reveals accelerated cooperative behavior but no competitive disadvantage in mixes with rhamnolipid-deficient free riders. **a.** Swarming start times with and without autoinducer in the plate media. No QS: 8 biological replicates with 80 total technical replicates. QS: 6 biological replicates with 54 total technical replicates. p-value <1e-8 by Kolmogorov-Smirnov test. **b.** WT PA14 was competed at a 1:1 ratio against the Δ*rhlA* strain with quorum signals in the plate media (see methods). After competition for 24 hours the plates were washed and the cells diluted and counted by CFU. The change in the proportion of the WT strain (Final ratio WT/(Total Cell Number) – Initial Ratio (0.5)) at the conclusion of the competition is shown. Initial frequencies were recorded and final ratios calculated with respect to these. Data includes three biological replicates with several technical replicates in each. See Supplementary Table 3 for initial ratios and final population sizes for each competition.

With the observations that quorum signal perturbation leads to increased per capita gene expression in immotile colonies and earlier onset of tendril formation, we asked if this perturbation could involve a fitness cost when the WT is in competition with an established defector mutant (Xavier et al. 2011; de Vargas Roditi et al. 2013). Surprisingly, we saw no competitive cost to the WT when quorum signals were added to the swarming plate media (Figure 6b). Since there is no visible growth cost, we conclude that—despite our attempts to perturb *rhlAB* expression—swarming cooperation remains robust to cheating.

## Discussion

Here, we used a combination of experimental and computational methods to advance our knowledge of *rhlAB* expression in a spatially-structured environment. We developed a novel high-throughput analysis, using fluorescence to track spatial-temporal bacterial growth and gene expression. The data produced showed that the gene expression in motile swarming *P. aeruginosa* colonies peaked at the tip of each tendril, a finding unexpected from our previous understanding of *rhlAB* expression (Figure 1). This phenotype emerged regardless of cell motility (Supplementary Figure 4). Further, we found that swarming tendrils, while expanding at a linear velocity, were able to maintain an exponential growth rate. This exponential growth rate is generated by a sustained growth rate throughout the tendril, not localized to the tendril edge as expected. This may be an example of navigated range expansion, following recent observations of growth dynamics in motile *E. coli* strains (Cremer et al, 2019).

To explore this unintuitive phenotype, non-motile colonies started from single cells provided a valuable model to study the communication between bacterial aggregates via diffusible compounds impacting gene expression and cooperative behavior. We found the immotile colony an underappreciated model that produced massive amounts of data to characterize growth and gene expression with spatial interaction (Figure 2). Even with classic microbiology assays we believe new layers of regulation to bacterial behavior can be quantified, perturbed and characterized that were not present in the equivalent liquid culture experiments.

Our analysis not only revealed that *P. aeruginosa* colonies can communicate across centimeter scale distances (Figure 4), it showed that colony communication through the integration of growth and quorum signal information is capable of generating complex regions of both positive and negative interactions between colonies that further vary with the configuration of the cell aggregates and scale with the colony’s growth rate (Figure 3). The kernels of interaction that we built from these data can be used to generate hypotheses of relevant length and growth timescales that may provide insight into the robustness of social interaction and cooperative phenotypes in natural bacterial communities.

Furthermore, the imaging infrastructure we have described allows for high throughput iteration between the immotile and motile systems. The volume of the data we were able to collect using immotile colonies allowed us to uncover a spatially-linked perturbability to cooperative gene expression whereby the colonies are able to express *rhlAB* at levels previously uncharacterized by liquid culture experiments (Figure 4). Surface-induced gene expression has been seen before in biofilm polymer production and cell shape changes have been observed in the transition from planktonic life to swarming (Davies and Geesey 1995; Sauer et al. 2002; Kuchma and O’Toole 2000; Sauer and Camper 2001; McCarter and Silverman n.d.; Davies et al. 1993; Harshey and Matsuyama 1994). However, these studies focus on the presence of gene expression in bacteria attached to surfaces or present in biofilms that wasn’t present in liquid. *rhlAB* expression occurs regardless of surface or liquid environment in *P. aeruginosa*. In this study, we uncovered a new, spatially-linked perturbability to *rhlAB* expression, giving a social degree of freedom to the control of this cooperative behavior that becomes possible when cells are on a surface or moving in a swarm.

While our immotile colony data indicated there was no growth cost under social perturbation (Figure 5), work in the motile swarms indicated that this increased gene expression pattern played a role in the fully motile system (Figure 6a). This gives us a unique opportunity to ask if this behavior could play a role in the competitive motile system (Figure 6). Finding no competitive disadvantage to this increased expression pattern, we conclude this new regime of gene expression falls under the metabolic prudence regulation structure. However, we note that quorum signal perturbation likely has systemic effects and that the impact we observed on the time to tendril formation in the swarms is likely dependent on more than rhamnolipid production alone.

To crystalize how *rhlAB* expression changes with both growth rate and social environment, we submit Figure 7 as a model for this system. Biomass (red) growth rate (black) and per capita gene expression (green) are representations of data previously presented and analyzed (Figure 1). The quorum signal curve (blue) is an approximation of the quorum signal field assuming a constant production rate with growth rate. The key elements to the integration of nutrient limitation and quorum signal concentration are as follows. When the cells are growing with a high growth rate, cells are more susceptible to perturbation by quorum signal (Figure 3c and Figure 4d). This means that when biomass is growing at a similar rate and with a similar amount of biomass, as can be seen in comparing the center of a swarm before tendril formation (Figure 7a) with the tip of the tendril at 12 hours (Figure 7b), the tip of the tendril, exposed to a higher quorum signal concentration, will express more *rhlAB* per capita. This also means that when exposed to the same quorum signal levels but experiencing a difference in growth rate, cells growing at a slower rate will express less *rhlAB* per capita. This can be seen by comparing the gene expression at the slow-growing swarm center at 12 hours (Figure 7b) with the same quorum signal concentration experienced by the tip of the swarm tendril at 20 hours (Figure 7c). Taken together, a metabolically prudent basis for gene expression with high perturbability at high growth rate could explain the emergent directionality of *rhlAB* per captia gene expression that we observe experimentally.

**Figure 7:**
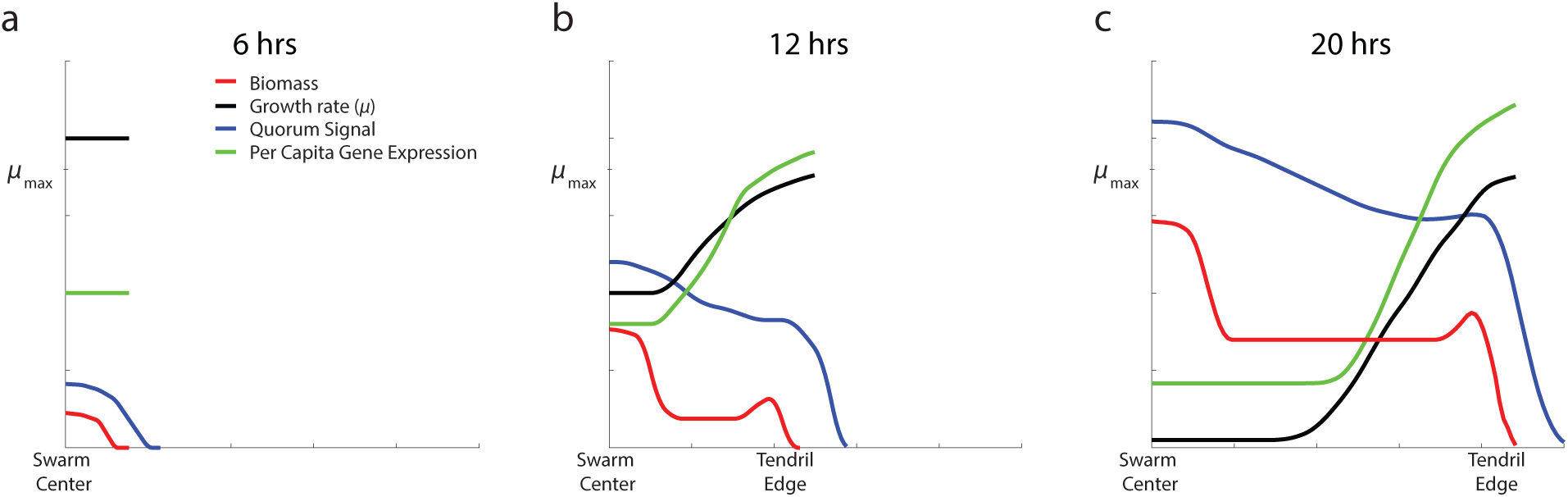
The combination of nutrient information and quorum signals allows for the emergence of directionality. Cartoon depicting three key timepoints in swarming tendril development. Cross sectional biomass density overlaid with distance dependent growth rate, quorum signal concentration and per capita *rhlAB* expression. **a**. At 6 hours, no tendrils have formed, biomass is largely localized at the original seeding location. Growth rates in the cells are high and quorum signals are less than maximal. Gene expression is uniform per unit of biomass throughout the population. **b**. At 12 hours, the tendril is moving at a constant velocity, growth is localized to the edge of the tendril and the biomass localizes at the tendril tip. Quorum signals continue to be produced in the center of the swarm, the rest of the tendril produces quorum signals proportional to the regions biomass level and growth rate. The gene expression that results is still moderate at the swarm center, but now highest at the tip of the tendril, correlating with the highest growth rate. **c**. At 20 hours, the tendril tip may be experiencing quorum signal levels similar to those of the center given a production rate proportional to the growth rate and diffusion of the signals. However, the comparatively high growth rate in this region allows a distinction between quorum signal levels accumulated over time (swarm center) and the resultant low gene expression, and quorum signal levels accumulated due to rapid growth and the corresponding high gene expression.

Bacteria exist in complex social and spatial environments, but very little is known regarding cellular decision-making in these highly dynamic and spatially-driven environments. The basis of our understanding of bacterial behavior comes from studying regulation mechanisms thoroughly but in liquid culture. However, bacteria live mostly in spatially-structured environments. Experimental models such as *P. aeruginosa* swarms and even immotile colonies growing on hard agar allow us to study proximate molecular mechanisms and ultimate evolutionary questions in spatially structured communities (Yan et al 2019). Spatially structured environments are able to recapitulate a range of behavior in natural bacterial communities unattainable by liquid culture experiments. By iterating experimental and computational data-driven methods we demonstrate that the integration of quorum signal and nutrient limitation information in the metabolically prudent regulation of *rhlAB* can still let the tip of a swarming tendril emerge as a region of high cooperative gene expression. Further, we show that this gene expression control is robust to cheating across a much wider range of conditions than previously appreciated by liquid culture experiments. As many social behaviors take in diffusive inputs, this result may be generalizable to a wide range of social or cooperative phenotypes with spatially-linked gene regulation that can already be assayed with classic microbiology techniques.

## Materials and Methods

### Microbiological assays

Plates were made with the recipe described in (Yan et al. 2019) with the addition of 1.8mL of L-arabinose at 40% weight/volume for a concentration of 1.5%. Water was subtracted to compensate. Every plate has 20mL of agar media and was inspected visually to confirm a flat surface. Swarms were prepared as described in (Yan et al. 2019). Liquid culture assays were performed using casamino acid media prepared as in (Xavier et al. 2011). Data was acquired on a benchtop TECAN plate reader. The data was analyzed using custom software in MATLAB (Boyle et al. 2015).

All timeseries were imaged with prototype imaging setup, Canary (Supplemental Figure 1). Fluorescent LEDs were used to light the sample. Data was collected by Atik VS14 Fluorescent Camera through the Thorlabs filter wheel FW102C. Timeseries was collected through a custom-built control system using the Arduino Uno R3. Immotile colony timeseries were imaged every 10 minutes. Swarms were imaged every 5 minutes. Images taken in Canary were subject to uneven lighting due to the placement of the fluorescence LEDs (Supplementary Figure 1). To correct for this, multiple plates were imaged in Canary as well as on a flatbed scanner. We called this scanner data our ‘ground truth’. A correction was built from these images that allowed us to take each image generated in Canary and alter it to the evenly lit environment on the scanner. This correction was built manually by extracting features from the images. The data was validated on rotational datasets taken in Canary (Supplementary Figure 3). The final background correction is shown below. Parameters vary depending on the exact configuration of Canary though the terms remain consistent. The correction was updated as the instrument received upgrades and to control for variation in the L-arabinose batch used.

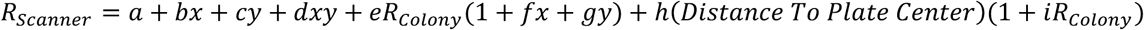

Min and max biomass and gene expression data full range boundaries from swarming tendrils (Figures 1bc) was smoothed once with a moving window of 5 for visualization. Background corrected pixel data is smoothed once with a moving window of 5 along the time axis before pixel data is extracted and grouped into colony components. Colony Red data is smoothed once with a moving window of 5 before exponential growth rates of the colonies are calculated.

Unless noted otherwise, swarms and colonies provided exogenous quorum signals were perturbed with the concentration of quorum signals determined to be present after 24 hours of swarming (Xavier et al. 2011).

### Analysis of immotile colonies

Cells were grown overnight in Casamino acid media and passaged into fresh Casamino acid media for 2-4 hours to reach exponential phase. These cells were then triple washed, diluted and spotted onto the agar such that every colony arises from a single cell. Colonies were plated with motility-preventing agar concentrations as in the classic Colony Forming Unit (CFU) assay (Figure 6a,b). The cells were fluorescently labeled for both biomass generation and rhamnolipid investment. Biomass was tracked using DsRed(DC2) (Pfleger et al. 2005) under the control of the PBad promoter induced by L-arabinose in the plate media (Figure 6c, Figure 2) (Newman and Fuqua 1999). Rhamnolipid investment was tracked through the previously validated P*rhlAB* -GFP promoter fusion (van Ditmarsch and Xavier 2011; Boyle et al. 2015).

The image timeseries post-processing was done in MATLAB R2018a (Supplemental Figures 3 and 5) and used to generate colony-centric growth, per captia gene expression information and all spatial-temporal features used in the text.

To supplement the identification of colonies, we developed a method to separate colonies that grow together and “merge” over the course of the timeseries so they could be tracked independently. After the images were background corrected, the peaks of the colonies were identified across a range of images and parameter values. The images used are between 20 and 30 hours, before the majority of colony merge events. The best parameters for peak identification were selected and used in the downstream analysis.

Each complete image timeseries was used to create a mask with all pixels that will eventually contain biomass. Once identified, each pixel was tracked throughout the timeseries. To localize pixels to their cognate colony, the previously identified peaks, the mask and the biomass distribution in the final timepoint were used with the watershed algorithm to identify the boundaries of colony objects.

L2 (Ridge) regularization was performed with a 4-fold cross validation (Figure 3c).

### Analysis of swarming colonies

To determine the speed of a moving tendril, the location of the edge every 15 minutes between 12 and 20 hours was calculated and the data was smoothed with a moving window of 1.25 hours. Swarms were imaged in a prototype imager equipped with a fish eye lens allowing for the acquisition of brightfield data for up to twelve swarming plates at a time. The timeseries were analyzed in ImageJ to identify the time of tendril formation. As the fish eye lens spreads the image pixels to cover a much larger region, the signal to noise ratio was managed carefully when collecting these data. Tendril formation times for each plate were calculated at three different zoom levels and averaged. To avoid bias, the data for each plate was collected by at least two independent researchers before averaging.

## Acknowledgements

The authors acknowledge Ned Wingreen, Chris Myers, Kyu Rhee, Dan Heller, Jinyuan Yan, Bradford P. Taylor and Chen Liao for helpful discussions and manuscript comments. This work was funded by National Science Foundation (www.nsf.gov) award MCB-1517002/NSF 13-520 to JBX and a National Science Foundation Graduate Research Fellowship GRFP DGE-1257284 2012 to HM.

