## Supplementary Materials for "Spatial-temporal dynamics of a microbial cooperative behavior robust to cheating"

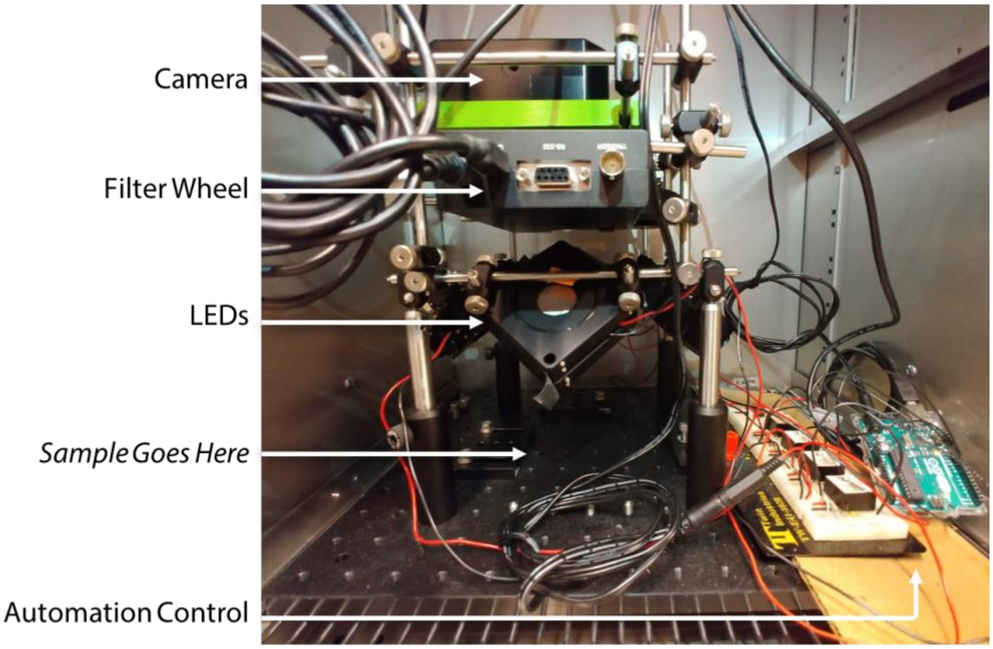


**Supplemental Figure 1: Canary diagram.** Canary is a custom-built imaging device designed to allow fluorescent imaging inside an incubator [See methods].

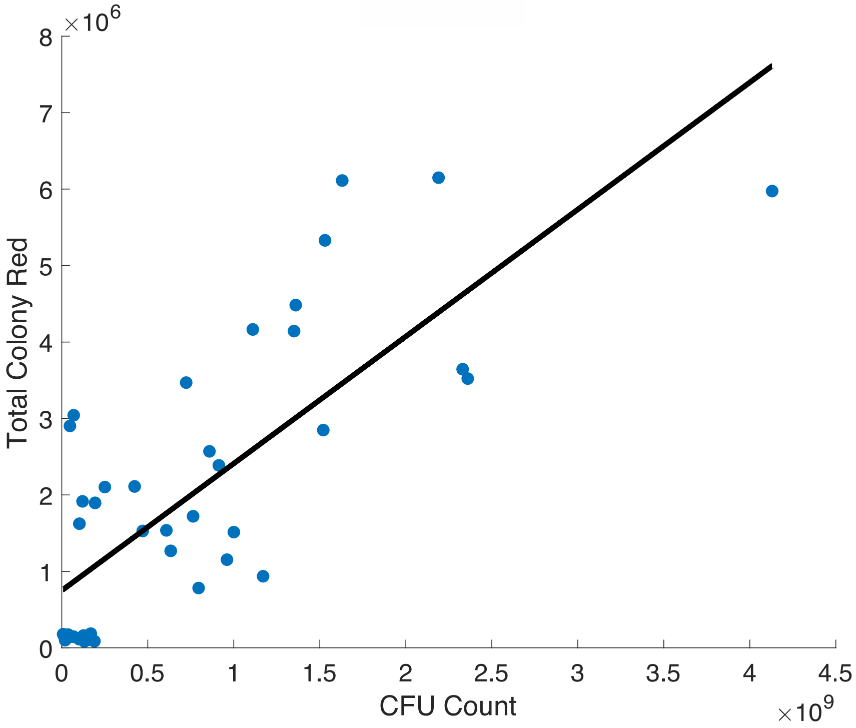


**Supplemental Figure 2: DsRed as a proxy for colony biomass.** The total fluorescence for the colonies picked was compared to the colony's CFU count. Fluorescence data was collected by scanning CFU plates on a flatbed plate scanner. Colonies were picked at various growth stages and assayed for total CFU. $R_2$ = 0.6006.

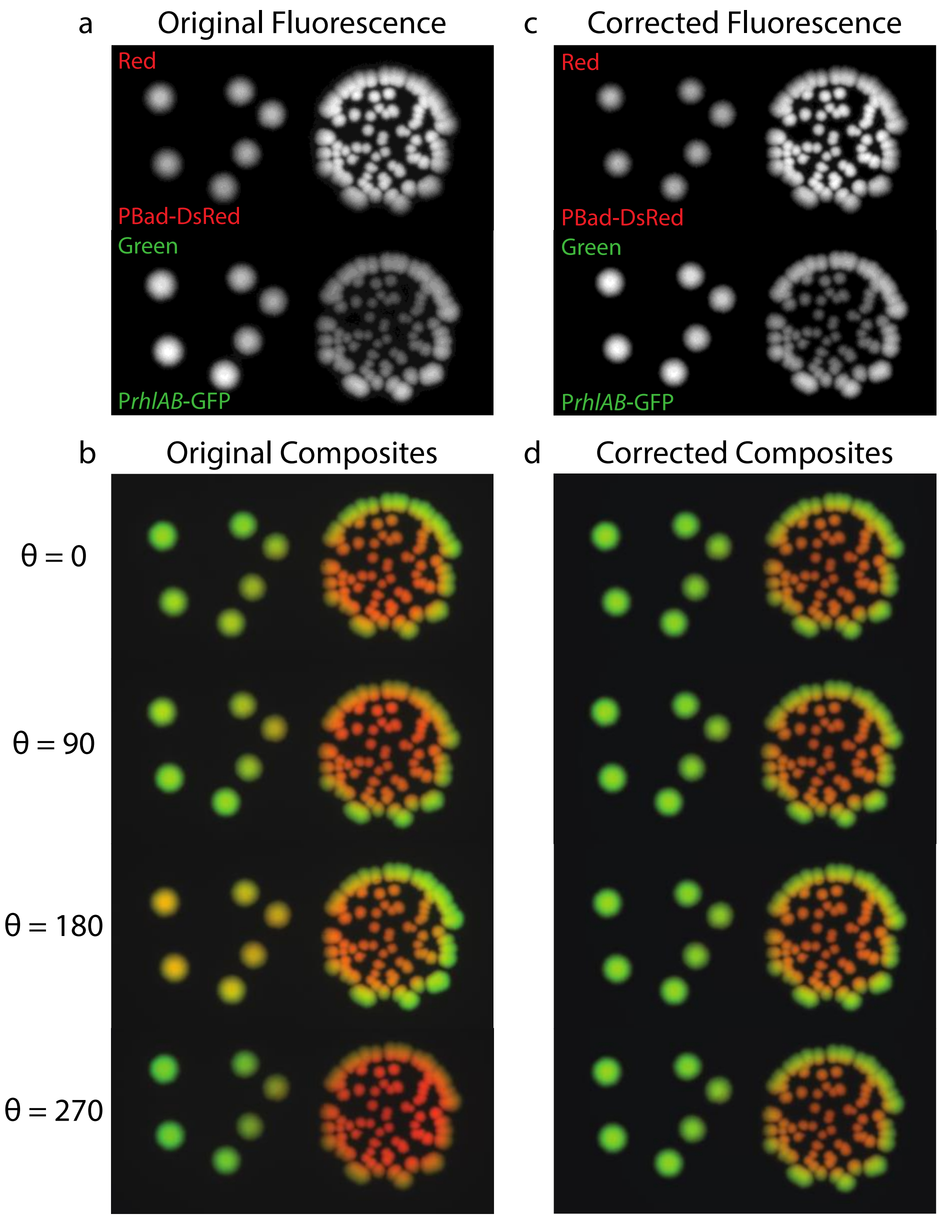


**Supplemental Figure 3: Background Correction. a.** 48 hour images before background correction [top] DsRed fluorescence [bottom] GFP fluorescence **b.** 48 hour RGB composite images before background correction **c.** RGB composite images after background correction (see Supplementary Methods Text (and equ below) for a detailed derivation of the correction. **d.** 48-hour images after background correction [top] DsRed fluorescence [bottom] GFP fluorescence

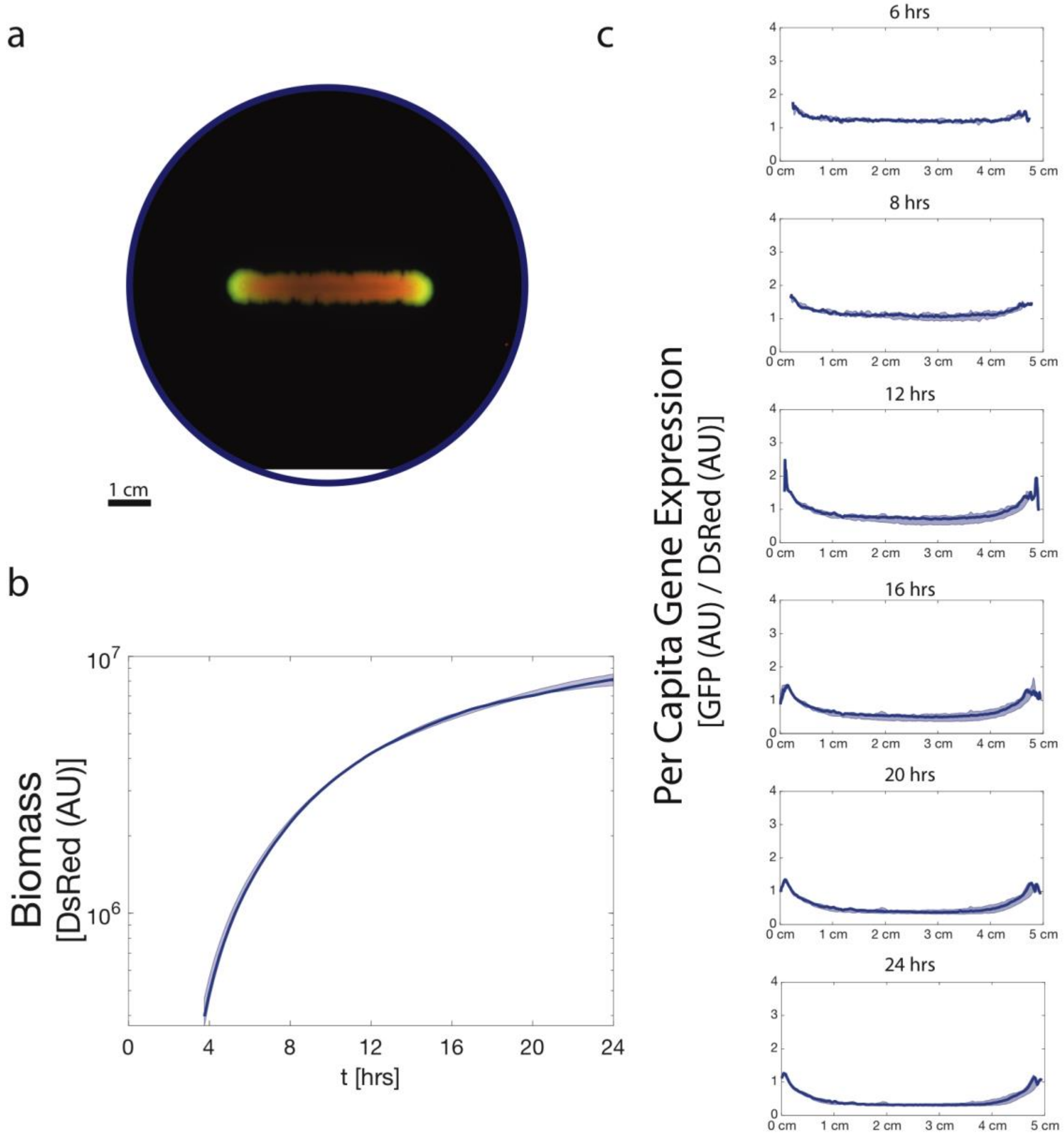


**Supplemental Figure 4: Immotile tendrils show growth saturation and increased per capita gene expression at the edges. a.** Image of an immotile tendril experiment at 48 hours. 10 $\mu$L of washed exponential phase culture at OD600 of 1 was seeded uniformly along a 5mm line and measured in timeseries. Labeling schema as previously described. **b.** Growth in the immotile tendril. In contrast to the swarming tendrils, the cells are unable to maintain an exponential growth rate. Shaded region describes full range of the data across three biological replicates. **c.** Per capita gene expression along the length of the tendril. The edge of the immotile tendril continues to emerge as regions of high expression here as in the swarming tendrils. Shaded region describes the full range of the data across three biological replicates.

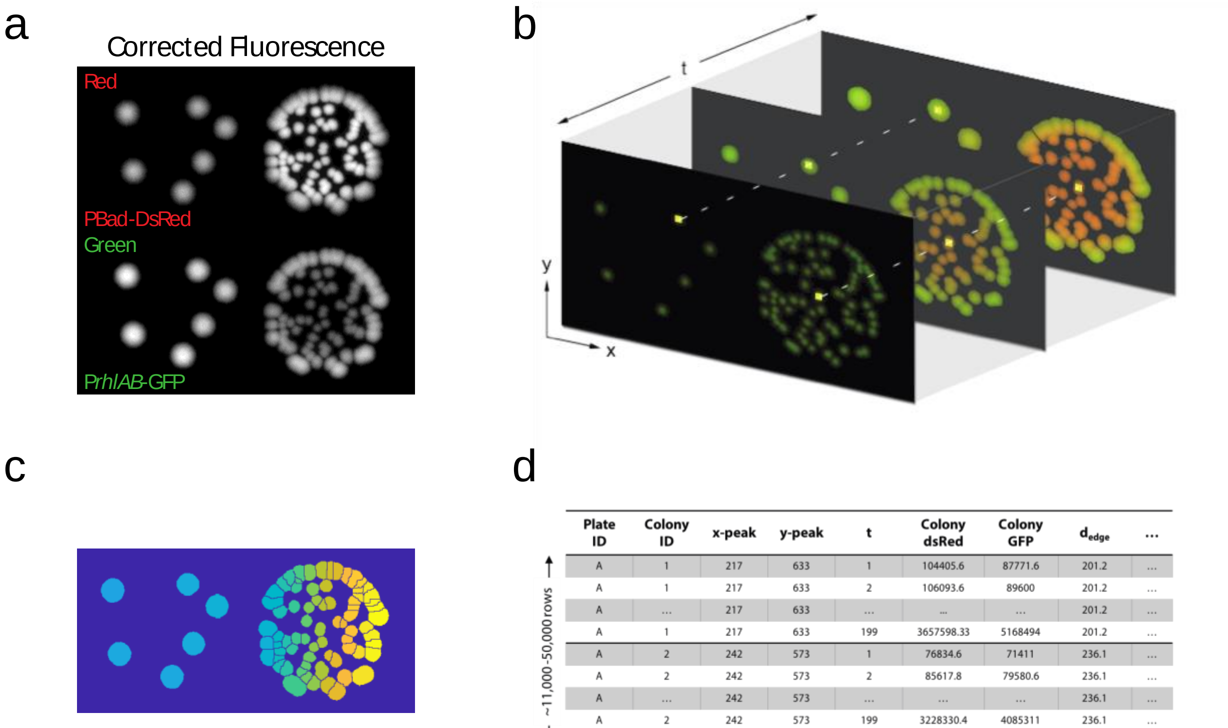


d

| Plate ID | Colony ID | x-peak | y-peak | t | Colony dsRed | Colony GFP | $d_{edge}$ | ... |
|---|---|---|---|---|---|---|---|---|
| A | 1 | 217 | 633 | 1 | 104405.6 | 87771.6 | 201.2 | ... |
| A | 1 | 217 | 633 | 2 | 106093.6 | 89600 | 201.2 | ... |
| A | ... | 217 | 633 | ... | ... | ... | 201.2 | ... |
| A | 1 | 217 | 633 | 199 | 3657598.33 | 5168494 | 201.2 | ... |
| A | 2 | 242 | 573 | 1 | 76834.6 | 71411 | 236.1 | ... |
| A | 2 | 242 | 573 | 2 | 85617.8 | 79580.6 | 236.1 | ... |
| A | ... | 242 | 573 | ... | ... | ... | 236.1 | ... |
| A | 2 | 242 | 573 | 199 | 3228330.4 | 4085311 | 236.1 | ... |
| ⋮ | ⋮ | ⋮ | ⋮ | ⋮ | ⋮ | ⋮ | ⋮ | ⋮ |

~11,000-50,000 rows

~400 columns

**Supplemental Figure 5: Pipeline Workflow. a.** CFU images after background correction **b.** Visualization of pixel-based approach for image timeseries data extraction. **c.** Pixels are collated using a custom image analysis algorithm within each colony and tracked across time. Final pixel to colony allocation map shown. **d.** Mockup of a pipeline output data frame.

**Supplementary Table 1: Goodness of fit for regularized regression models**

| Test | $R_2$ |
|---|---|
| $0 \leq \mu < 0.25$ | 0.2993 |
| $0.25 \leq \mu < 0.5$ | 0.6066 |
| $\mu \geq 0.5$ | 0.6374 |

**Supplementary Table 2: Mann-Whitney Test Results to Figure 6**

| Test | WT No QS vs WT +QS | WT +QS vs WT +QS Config Control | WT No QS vs WT +QS Config Control |
|---|---|---|---|
| Growth Rate | 0.0689 | 0.0599 | 0.6419 |
| Final Biomass | 0.0435 | <1e-10 | <1e-10 |
| Time of Colony Appearance | 0.0014 | <1e-10 | <1e-10 |

**Supplementary Table 3:**

| Biological Replicate | WT Initial Proportion (WT/Total) | Competition Change in Ratio (Final - Initial WT Proportion) | Final Population Size |
|---|---|---|---|
| 1 | 0.55 | 0.04 | 6.74E+08 |
| 1 | 0.55 | 0.15 | 7.18E+08 |
| 1 | 0.55 | 0.03 | 1.72E+09 |
| 1 | 0.55 | 0.08 | 1.61E+09 |
| 1 | 0.55 | 0.14 | 7.79E+08 |
| 1 | 0.55 | 0.03 | 5.51E+08 |
| 1 | 0.55 | 0.06 | 6.39E+08 |
| 1 | 0.55 | 0.07 | 3.68E+08 |
| 1 | 0.55 | 0.14 | 5.60E+08 |
| 1 | 0.55 | 0.23 | 2.71E+08 |
| 1 | 0.55 | 0.10 | 1.23E+08 |
| 1 | 0.55 | 0.12 | 2.10E+08 |
| 2 | 0.54 | 0.10 | 7.16E+09 |
| 2 | 0.54 | 0.14 | 4.50E+09 |
| 2 | 0.54 | 0.09 | 3.71E+09 |
| 2 | 0.54 | 0.10 | 7.93E+09 |
| 2 | 0.54 | 0.11 | 8.94E+09 |
| 2 | 0.54 | 0.13 | 1.69E+10 |
| 2 | 0.54 | 0.11 | 1.64E+10 |
| 2 | 0.54 | 0.09 | 3.52E+09 |
| 2 | 0.54 | 0.12 | 6.62E+09 |
| 2 | 0.54 | 0.11 | 8.56E+09 |
| 2 | 0.54 | 0.14 | 7.11E+09 |
| 2 | 0.54 | 0.10 | 1.31E+10 |
| 3 | 0.49 | 0.05 | 2.43E+09 |
| 3 | 0.49 | 0.14 | 4.87E+09 |
| 3 | 0.49 | 0.09 | 1.56E+10 |
| 3 | 0.49 | 0.14 | 2.94E+09 |
| 3 | 0.49 | 0.12 | 5.83E+09 |
| 3 | 0.49 | 0.08 | 7.26E+09 |
| 3 | 0.49 | 0.01 | 8.59E+09 |
| 3 | 0.49 | 0.12 | 7.28E+09 |
| 3 | 0.49 | 0.10 | 1.27E+10 |
| 3 | 0.49 | 0.12 | 1.06E+10 |
| 3 | 0.49 | 0.11 | 4.59E+09 |
